# Neuro-functional correlates of protective effects of wheel-running exercise against cocaine locomotor sensitization in mice: a [^18^F]fallypride microPET study

**DOI:** 10.1101/855791

**Authors:** Guillaume Becker, Louis-Ferdinand Lespine, Mohamed Ali Bahri, Maria Elisa Serrano, Christian Lemaire, André Luxen, Ezio Tirelli, Alain Plenevaux

**Author notes:** These auhtors contributed equally to this work (Guillaume Becker, Louis-Ferdinand Lespine). Corresponding author: Guillaume Becker, Radiobiology Unit, SCK•CEN, Belgian Nuclear Research Centre., Boeretang 200, 2400Mol, Belgium., GIGA – Cyclotron Research Center – In Vivo Imaging, University of Liège., Quartier Agora – B30, Allée du 6 Août, 8., 4000 Liège, Belgium.

## Abstract

Wheel-running exercise in laboratory rodents (animal model useful to study the neurobiology of aerobic exercise) decreases behavioral markers of vulnerability to addictive properties of various drugs of abuse including cocaine. However, neurobiological mechanisms underpinning this protective effect are far from being fully characterized and understood. Here, 28-day-old female C57BL/6J mice were housed with (n=48) or without (n=48) a running wheel for 6 weeks before being tested for acute locomotor responsiveness and initiation of locomotor sensitization to intraperitoneal injections of 8 mg/kg cocaine. The long-term expression of sensitization took place 3 weeks after the last session. On the day after, all mice underwent a microPET imaging session with [^18^F]fallypride radiotracer (dopamine 2/3 receptor (D2/3R) antagonist). Exercised mice were less sensitive to acute and sensitized cocaine hyperlocomotor effects, such attenuation being particularly well-marked for long-term expression of sensitization (*η*^*2*^*p* = 0.262). Additionally, we found that chronic administrations of cocaine was associated with a clear-cut increase of [^18^F]fallypride binding potential in mouse striatum (*η*^*2*^*p* = 0.170), presumably reflecting an increase in postsynaptic D2/3R density in this region. Finally, we found evidence that wheel-running exercise was associated with a moderate decrease in D2/3R density in striatum (*η*^*2*^*p* = 0.075), a mechanism that might contribute to protective properties of such form of exercise against drugs of abuse vulnerability.

## INTRODUCTION

By the end of the 90’s, society’s perspective on addiction changed, promoted in a certain extent by a new pathophysiological vision of addictive disorders, dubbed as the Brain Model Disease of Addiction^1^. Although the concept of addiction as a brain’s disease is still being questioned^2,3^, in vivo neuro-imaging has greatly impacted our pathophysiological understanding of the chronic brain state of addiction^4–8^. Imaging studies have evidenced a decreased dopamine D2 receptors availability in the dorsal striatum of patients suffering from cocaine addiction^9,10^. Preclinical imaging investigations on non-human primates have shown similar results following cocaine self-administration^11,12^.

The actual trend in public healthcare policies combating drug abuse is to promote prevention strategies to reduce the risks of substance addiction development^13^. In this context, physical exercise has been promoted as its benefits are supported by cross-sectional and longitudinal studies conducted on adolescents and young adults reporting negative association between physical exercise or sports participation and the initiation of drugs of abuse consumption^14–16^. Additionally, physical activity has been shown to exert curative effects by attenuating relapse rates in alcohol, nicotine and illicit drugs abusers^15,17^. However, the neurobiological mechanisms that underlie this relationship are still needed to be uncovered. Preclinical studies using animal models useful for the study of addiction have provided evidence for both preventive-like and curative-like effects of physical activity on drugs of abuse vulnerability. In rodents studies, physical exercise is often modelled by the use of a freely available running wheel place in the housing cages. With such a paradigm, exercised rats exhibited reduced rates of acquisition, motivation or escalation of self-administration of cocaine, heroin, methamphetamine or speedball, as compared to sedentary animals^18–23^. Consistent with self-administration reports, wheel-running exercise has also been shown to be effective at reducing the acute and chronic locomotor-stimulating effects of cocaine as well as the expression of sensitization to those effects^24,25^. This phenomenon, associated with major neurochemical changes in dopaminergic and glutamatergic systems notably^26^, is thought to play an integral role in craving and relapse^27,28^.

Behavioral sensitization (e.g. locomotor sensitization) has been define as a progressive and enduring augmentation in the locomotor activating and reinforcing effect of a psychostimulant, consecutively to repeated exposure to these drugs. This phenomenon, and the underlying neuropathological processes, has been thought to be useful for studying the neuronal adaptations that leads to compulsive drug craving related to the induction and expression of sensitization.

As a consequence, one of the most challenging issue in the biology of addiction rises from preclinical evidences consistently reporting that psychostimulant-treated rodents (cocaine and amphetamine), after a withdrawing period, are hypersensitive to the psychomotor activating and incentive motivational effects of these drugs^29^, More strikingly, in such sensitization experimental protocols, rodents are sensitized to the psychomotor effects of direct-acting D2 agonists^30,31^. In rodents, the effect of repeated exposure to cocaine on the D2 receptor availability remains unclear, with reports of increases^32,33^, decreases^34^ or unchanged^35,36^. Some of these discrepancies may be due to differences in experimental designs and protocols (*e*.*g*. doses and routes of injection, duration of drug withdrawal and timing of the expression of sensitization), as well as lack of statistical power^37^. It has been hypothesized that chronic amphetamine or cocaine administration could results in an increase in D2 high affinity state receptors density, whereas the total amount of receptors (low and high affinity states) may be unchanged^38,39^. These phenomenon, also described for nicotine chronic administration, may explain the hypersensitivity of psychostimulant-treated rodents^40^.

In our previous work, we reported in C57Bl6 mice that the effectiveness of wheel-running exercise at attenuating cocaine locomotor sensitization not only resisted to exercise cessation but was also unambiguously persistent^41^. Further on, we found that early-life period such as early adolescence (vs early adulthood) may be particularly sensitive to protective properties of this form of exercise against vulnerability to cocaine-induced locomotor sensitization^42^.

The purpose of the present study was twofold. First we aimed to replicate our previous behavioral results in female C57Bl6 mice. Second, we aimed to investigate the neuro-functional correlates of protective properties of wheel-running exercise on cocaine locomotor sensitization by assessing dopamine D2/3 receptors (D2/3R) availability with [^18^F]fallypride microPET. We ambitioned to test whether the positive effect of exercise was linked to a decreased D2/3R availability.

## METHODS

### Subjects

Ninety-six 21-day-old females C57BL/6J mice were obtained from JANVIER, Le-Genest-Saint-Isle, France. The choice of C57BL/6J strain was based on its extensive use in addiction research and previous experiments performed in our laboratory. Given available resources, we did not investigate sex-related difference in the interaction between exercise and cocaine responsiveness in favor of statistical power (i.e. higher sample size). We tested female as they may receive more benefits from exercise than males^42–44^. Upon arrival, mice were housed in groups of eight in large transparent polycarbonate cages (38.2 × 22 cm surface × 15 cm height; TECHNIPLAST, Milano, Italy) for one week of acclimation. On the following day, they were housed individually according to the experimental housing conditions (exercise or sedentary receiving cocaine or saline during testing, see section “Experimental Design and Procedure”) in smaller TECHNIPLAST transparent polycarbonate cages (32.5 × 17 cm surface × 14 cm height) with pine sawdust bedding, between-animal visual, olfactory and acoustic interactions remaining possible. Tap water and food (standard pellets, CARFIL QUALITY, Oud-Turnhout, Belgium) were continuously available. The animal room was maintained on a 12:12 h light-dark cycle (lights on at 07.00 a.m.) and at an ambient temperature of 20-23°C. All experimental treatments and animal maintenance were reviewed by the University of Liège Animal Care and Experimentation Committee (animal subjects review board), which gave its approval according to the Belgian implementation of the animal welfare guidelines laid down by the European Union (“Arrêté Royal relatif à la protection des animaux d’expérience” released on 23 May 2013, and “Directive 2010/63/EU of the European Parliament and of the Council of 22 September 2010 on the protection of animals used for scientific purposes”). All efforts were made to minimize the number of animals used and their suffering. Moreover, the ARRIVE guidelines (Animal Research Reporting In Vivo Experiments), which have been developed to improve quality of experimenting and reporting in animals studies, were followed as closely as possible^45^.

### Aerobic Voluntary Exercise

A running wheel was made of an orange polycarbonate saucer-shaped disk (diameter 15 cm, circumference 37.8 cm; allowing an open running surface) mounted (bearing pin) on a plastic cup-shaped base (height 4.5 cm) and tilted at a 35° angle from the vertical plane (ENV-044, Med Associates; St Albans, VT, USA). The base was fixed on a stable transparent acryl-glass plate. Running was monitored and recorded continuously during the 42-day pre-testing period via a wireless system, each wheel being connected to a USB interface hub (DIG-804, Med Associates) which relayed data to a Wheel Manager Software (SOF-860, Med Associates). Data dealing with wheel-running activity are shown in Fig. 1.

**Figure 1.**
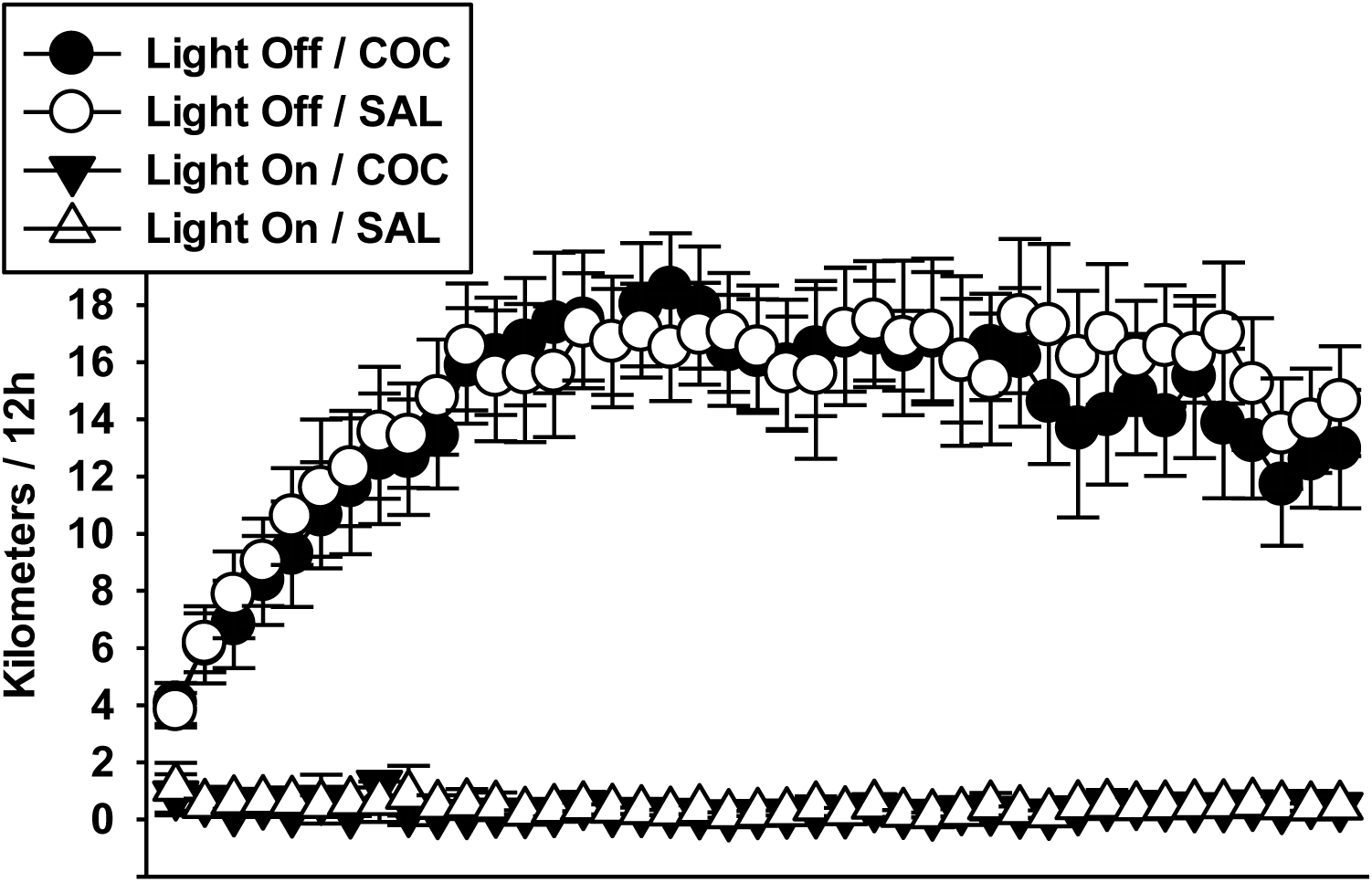
Wheel-running activity recorded prior to the testing period. Nocturnal (light off) and diurnal (light on) wheel-running activity of mice randomly assigned to exercise conditions. Since at this stage of the experiment mice from the cocaine (COC, n=24) and saline (SAL, n=24) groups were still undistinguishable, no inferential statistics were conducted on these data. All mice showed a rapid increase in wheel-running over the two first weeks until reaching a plateau. Bars represent 95% confidence intervals.

### Drug Treatments

(−)-Cocaine hydrochloride (BELGOPIA, Louvain-La-Neuve, Belgium), dissolved in an isotonic saline solution (0.9% NaCl), was injected intraperitoneally at a dose of 8 mg/kg in a volume of 0.01 ml/g of body weight, the control treatment consisting of an equal volume of isotonic saline solution. The dose and route of administration were selected on the basis of our previous studies^41,42^, these parameters being known to also induce rewarding-like effects in mice as measured by conditioned place preference^46^.

### Behavioral Test Chambers

A battery of eight chambers, connected to a custom written software for data collection, was used to measure mice locomotor activity, one mouse being tested in each chamber. Each activity chamber was constituted of a removable transparent polycarbonate tub (22 × 12 cm surface × 12 cm height), embedded onto a black-paint wooden plank serving as a stable base. The lid was made of a transparent perforated acryl-glass tablet. Two photocell sources and detectors were mounted on the plank such that infrared light-beams were located on the two long sides of the tub at 2-cm heights from the floor, 8-cm apart and spaced 6.5 cm from each end of the tub. Locomotor activity was measured in terms of crossings detected by the beams, one crossing count being recorded every time an ambulating mouse broke successively the two parallel beams. The activity chambers were individually encased in sound-attenuated shells that were artificially ventilated and illuminated by a white light bulb during testing. Each shell door comprised a one-way window allowing periodic surveillance during testing.

### [^18^F]fallypride radiosynthesis

The radiotracer [^18^F]fallypride was synthetized according to a method previously reported by Brichard *et al*. with slight modifications^47^. Briefly, the no-carrier-added synthesis of [^18^F]fallypride was conducted by nucleophilic substitution with [^18^F]fluoride of the p-toluenesulfonyl group of the commercially available precursor (ABX, Advanced Biochemical Compounds, Radeberg, Germany). After the labelling reaction that was conducted in acetonitrile (1 mL) with 3.5 mg of the substrate at 120 °C for 5 min, the crude reaction mixture was diluted with water (6 mL) and the resulting solution injected on a semi-preparative HPLC column. The purification was carried out at 254 nm using a Phenomenex Luna C18 column (5 μm, 250 × 15 mm) at a flow rate of 7 mL/min with an isocratic eluent of water/acetonitrile/trietylamine (45:55:0.1%; retention time of 22 min). The subsequent formulation step^48^ was realized by passing the HPLC collection solution, previously diluted with sodium chloride 0.9% (30 mL) and sodium ascorbate (30 mg) through a tC18 cartridge (360 mg, Waters). [^18^F]fallypride was then eluted from the support with ethanol (1 mL) and diluted with an isotonic solution (6 mL) containing sodium ascorbate (10 mg) as stabilizer. Based on the starting activity recovered from the cyclotron (111 GBq), this process afforded batches of [^18^F]fallypride ready for subsequent dilution for animal injection. The radiochemical yield was of 32 ± 5% (mean ± SD, decay corrected; n=24). At the end of beam, the averaged specific activity was 49 ± 20.4 Ci/µmol (1813.6 ± 755.6 GBq/µmol, decay corrected, n=24) and the synthesis duration of about 50 min. All the process was automated on a FASTlab synthesizer from GE Healthcare with single use components.

### [^18^F]fallypride microPET imaging data acquisition and processing

Twenty-four [^18^F]fallypride microPET imaging sessions were completed and all necessary efforts were made to systematically repeat the same procedure. Anesthesia was induced with 4% of isoflurane, afterward the mice were placed prone in a dedicated bed. Anesthesia was maintained with 1–2% isoflurane in a mixture of air and oxygen (30%) at 0.6 l/min. A stereotaxic holder (Minerve, Esternay, France) was systematically used to reduce head movements. Respiratory rate and rectal temperature were permanently measured using a physiological monitoring system (Minerve, Esternay, France). Temperature was maintained at 37 ± 0.5° C, using an air warming system.

[^18^F]fallypride was administered as bolus intravenous injection in the lateral tail vein over 20 seconds with a mean injected activity of 12.4 ± 3 MBq (range: 4.9 – 19.1 MBq). The mean injected mass of fallypride was 0.29 ± 0.35 µg (range: 0.02 – 2.51 µg). At the time of injection, dynamic microPET scans over 60 minutes were acquired in list-mode using a Siemens Concorde Focus 120 microPET (Siemens, Munich, Germany) and followed by 10 minutes transmission measurement with ^57^Co point source. The list-mode emission data were histogrammed into three-dimensional (3D) sinograms by Fourier rebinning and reconstructed by filtered backprojection with a ramp filter cutoff at the Nyquist frequency. All Corrections were applied except for scatter events^49^. No partial volume correction was performed on the acquired data. A set of 3D images was reconstructed in a 256 × 256 × 95 matrix and a zoom factor of 2. The reconstructed voxel size was 0.4 × 0.4 × 0.8 mm^3^. The dynamic time framing was set as follows: 6 × 5 s, 6 × 10 s, 3 × 20 s, 5 × 30 s, 5 × 60 s, 8 × 150 s, 6 × 300 s, and all data were decay corrected to the beginning of each individual frame.

Immediately after PET acquisition, the anesthetized mice were transferred into a 9.4 Tesla MRI DirectDrive VNMRS horizontal bore system with a shielded gradient system (Agilent Technologies, Palo Alto, CA, USA). A 72-mm inner diameter volumetric coil and a 2-channels head surface coil (Rapid Biomedical GmbH, Würzurg, Germany) were used as transmitter and receiver coils, respectively. The 3D anatomical T2-weighted brain images were acquired with a fast spin echo multislice sequence using the following parameters: TR/TEeff = 2500/40 ms, matrix = 128 × 128 × 64, FOV = 20 × 20 × 10.5 mm^3^, voxel size: 0.156 × 0.156 × 0.164 mm^3^, and a total acquisition time of 21 min.

Imaging data were processed with PMOD software (version 3.7, PMOD Technologies Ltd., Zurich, Switzerland). The processing includes a manual rigid co-registration of individual MRI images to its corresponding PET images, a spatial normalization of the co-registered MRI onto the PMOD MRI template, and the extraction of the PET time-activity curves of the left and right striatum as well as the cerebellum. Briefly, the inverse deformations parameters obtained during the spatial normalization of the individual MRI images onto the PMOD template were used to bring the mouse brain atlas^50,51^ into the native dynamic PET space and then extract the TACs based on the atlas predefined structures. The extracted TACs were then transferred into the kinetic modelling module of PMOD in order to estimate the [^18^F]fallypride binding. The non-displaceable binding potential (BP_ND_) parameter^52^ was calculated using the multi-linear reference tissue model (MRTM2) with the cerebellum as reference tissue^53^. We controlled the homogeneity of the TACs in the reference region (i.e. cerebellum) between each group, to rule out any bias from radiotracer inputs variations in the reference region. The statistical analysis revealed no difference between the four groups of the study for the cerebellum [^18^F]fallypride TACs expressed as area under the curve (Supp.1).

### Experimental Design and Procedure

Experimental timeline and design are presented in Fig.2. Ninety-six mice were housed in exercised (n=48) or sedentary (n=48) conditions from 28 days of age (early adolescence; ref needed) and were kept in these conditions until the end of behavioral experimentation. Since mice from the two housing environments received cocaine or saline during testing, a basic 2 (housing conditions: EX vs SED) × 2 (pharmacological treatment: COC vs SAL) factorial design was generated with N=96, n=24 per group based on preliminary results indicating increasing and decreasing effects of cocaine (vs saline, *η*^*2*^*p* = 0.162) and exercise (vs sedentary, *η*^*2*^*p* = 0.093) respectively on [^18^F]fallypride BP_ND_. Note that experimental procedures and parameters associated with psychopharmacological tests were similar to those used in Lespine and Tirelli^42^ where continuously exercised females C57BL6/J were found to be less vulnerable than their sedentary counterparts to acute and sensitized locomotor responsiveness to cocaine. Testing included the following five phases. (1) A pre-test habituation session to familiarize animals to novelty of the test context without neither injection nor measurements. (2) A 2^nd^ drug-free session evaluating baseline locomotor activity under saline. (3) Nine once-daily injections of cocaine or saline, with the measurement of locomotor-activating effect of cocaine after each injection, initiating locomotor sensitization after the baseline session. (4) Taking place 21-23 days after the last cocaine injection, a session assessing expression of sensitization on which animals received their previous respective pharmacological treatment. Throughout psychopharmacological testing, mice were weighed and received their pharmacological treatment right before being placed in the test chamber, recording of crossings lasting 30 min in all sessions. Experimental blinding was not realized because the unique experimenter inevitably knew the housing condition and the pharmacological treatment of each mouse. (5) Twenty-four hours after the test of expression of sensitization, mice underwent [^18^F]fallypride microPET scan. Note that the wheels (for exercised mice) were removed 24h before neuroimaging measurement to avoid any potential effect of overnight wheel-running exercise on neuro-functional measures.

**Figure 2.**
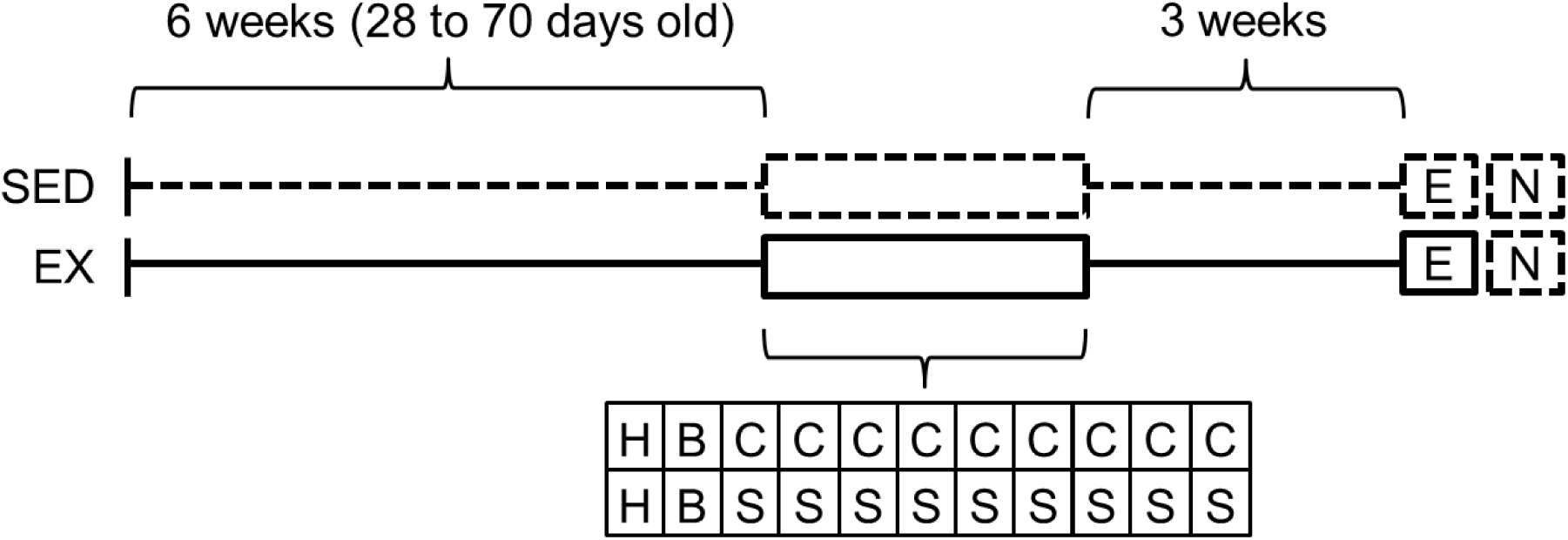
Experimental timeline and design. At 28 days of age, 96 mice were housed individually either in the presence (EX, n=48) or the absence (SED, n=48) of a running wheel. Testing began after 6 weeks in these housing conditions (from 28 to 70 days old). The experiment comprised four groups, mice from each housing group (EX or SED) receiving either cocaine or saline (with n=24 per group). Solid lines represent the presence of a running wheel in the home-cage and dotted lines its absence. H: habituation session (to familiarize animals to the novelty of the test context without neither injection nor measures); B: baseline session; the 2^nd^ once-daily session assessing the baseline activity under saline; C: cocaine intraperitoneal administration (9 once-daily sessions); S: control animals receiving saline intraperitoneal administration (9 once-daily sessions). E: session on which the expression of the sensitization was assessed 21-23 days after the last sensitizing injection, and under the previous pharmacological treatments. N: neuro-functional measures 24 h after the test of expression (microPET).

Due to practical reasons, the whole experiment was organized into twelve lots purchased and tested successively (each lot consisting of 8 mice). In each lot, two mice were assigned to one of the four experimental group by means of a computer-generated randomization schedule, the eight mice housed in acclimation cages contributing to theses four possible groups (sedentary/cocaine, sedentary/saline, exercised/cocaine, and exercised/saline). Therefore, the four groups were systematically represented within each lot by 2 mice to take into account any between-lot variability as well as that due to time and circumstances of testing (i.e. randomized block design, Supp.2). Additionally, due to impracticality to test 8 mice in a row on the same micro-PET scanning session, each block (n=8) was further split into 2 blocks (n=4) for the test for expression of sensitization and microPET imaging procedures. Again, the four groups were systematically represented within each block by one mouse (Supp.2). Therefore, mice were tested for expression of sensitization either 21 (half) or 23 (other half) days after the last cocaine injection, while all mice underwent neuroimaging scan 24h after this test. Note that the order of neuroimaging session was counterbalanced across subjects to avoid potential bias due to the specific activity variations. Experimenters conducting neuroimaging testing and analysis were blinded to experimental groups.

### Data Analysis

Inferential statistics were computed on the following data. (1) Acute responsiveness to locomotor-activating effects of cocaine scored as the difference between values derived from the first cocaine session and those of the baseline session. (2) Overall responsiveness to locomotor-activating effects of cocaine over the initiation of sensitization (9 sessions) scored as the area under the curve with respect to zero (AUC ground; calculation formula are based on and detailed by Pruessner and coll.^54^). (3) Locomotor activity exhibited during the expression of sensitization. (4) The bilateral [^18^F]fallypride BP_ND_ from left and right striatum of each subject were averaged to give a single [^18^F]fallypride BP_ND_ value per subject.

Each set of data was treated according to a randomized block design with a fixed-model 2 × 2 ANOVA incorporating the housing condition (EX or SED; 2 levels) and pharmacological treatment (COC or SAL; 2 levels) as between-group factors, and with the lot as a blocking factor (with 12 or 24 levels for the behavioral and neuro-functional measures respectively, see Supplement 1) followed by planned crossed or simple contrasts^55^. Each contrast was derived from the mean-square error term (MSE) provided by the ANOVA. Based on previous experiments or preliminary results, exercised mice were expected to display (1) lower cocaine locomotor responsiveness than sedentary mice (crossed contrasts) and (2) lower values of BP_ND_ (simple contrast). Cocaine-receiving mice were expected to show (1) greater locomotor activity and (2) higher BP_ND_ values than control saline mice (simple contrasts). Correlations were also computed to (1) determine whether the amount of wheel-running displayed before testing was associated with cocaine behavioral or neuro-functional outcomes, and (2) determine whether behavioral cocaine outcomes were associated with neuro-functional measures. Nocturnal distances over the 42-day pre-testing period were averaged for each (exercised) mouse, the resulting individual value serving as the measure of the overall distance travelled on the wheel. Effect sizes were given by *η*^*2*^*p*, Pearson correlation coefficient *r*, and probability of superiority (PS) where appropriate^56^. Statistical significance threshold was set at 0.05.

## RESULTS

### Psychopharmacological Measures

Fig. 3 Panel A depicts results dealing with baseline locomotor activity and initiation of cocaine locomotor sensitization over 9 once-daily sessions in mice housed either with or without a running wheel during adolescence (from 28 days of age) and tested 6 weeks later from 71 days of age. Panel B presents scores of acute responsiveness. Planned contrasts indicate that acute locomotor responsiveness to the 1^st^ administration of cocaine in sedentary animals was moderately attenuated in exercised mice (*η*^*2*^*p = 0*.*073, t*_(81)_ = 2.51, *p* = 0.007). As secondary outcomes, cocaine locomotor effect (vs saline, i.e. hyperlocomotor effect) was strong in each housing group (*t*s_(81)_ = 3.85 and 7.39, with a probability of superiority (PS) of 72 and 88 in exercised and sedentary groups respectively). In other words, the score of acute responsiveness from a cocaine-receiving mouse would be higher than that from a saline-receiving mouse for 72% of random pairs in exercised animals, and for 88% of random pairs in sedentary animals. Panel C depicts overall responsiveness during the initiation of sensitization scored as AUC ground. The pattern of results was comparable to that found for acute responsiveness with a more pronounced effect (*η*^*2*^*p* = 0.162, *t*_(81)_ = 3.96, *p* < 0.001). Cocaine effect was clear-cut in each housing group (*t*s_(81)_ = 5.47 and 11.07 with PS of 80 and 96 in exercised and sedentary groups respectively). Fig. 4 presents locomotor activity on the last (9^th^) session of sensitization (panel A, descriptive statistics) and on the test for expression of sensitization (panels B and C). Consistent with previous experimental stages, long-term expression of the sensitized locomotor responsiveness was largely reduced in exercised mice (*η*^*2*^*p* = 0.262, *t*_(81)_ = 5.37, *p* < 0.001). Again, cocaine effect was unambiguous in each housing group (*t*s_(81)_ = 5.48 and 13.08 with PS of 80 and 98 in exercised and sedentary groups respectively). Table 1 reports relationships between amounts of exercise and behavioral outcomes and [^18^F]fallypride BP_ND_, with correlation coefficients *r*s varying from −0.03 to 0.43 (*p*-values ranging from 0.89 to 0.035). Wheel-running exercise was strongly and positively associated with AUC ground in cocaine-receiving mice (*r* = 0.43). However, we think that this result should be (cautiously) interpreted in the context of other correlations results, number of tests performed, and sample size. Importantly, AUC ground was strongly and positively associated with expression of sensitization (found here at *r* = 0.55). The fact that wheel-running distances strongly correlate to AUC ground yet weakly to expression of sensitization (found here at r = 0.15), questions the nature of relationships reported between exercise and AUC ground (*i*.*e*. risk of false-positive).

**Fig 3.**
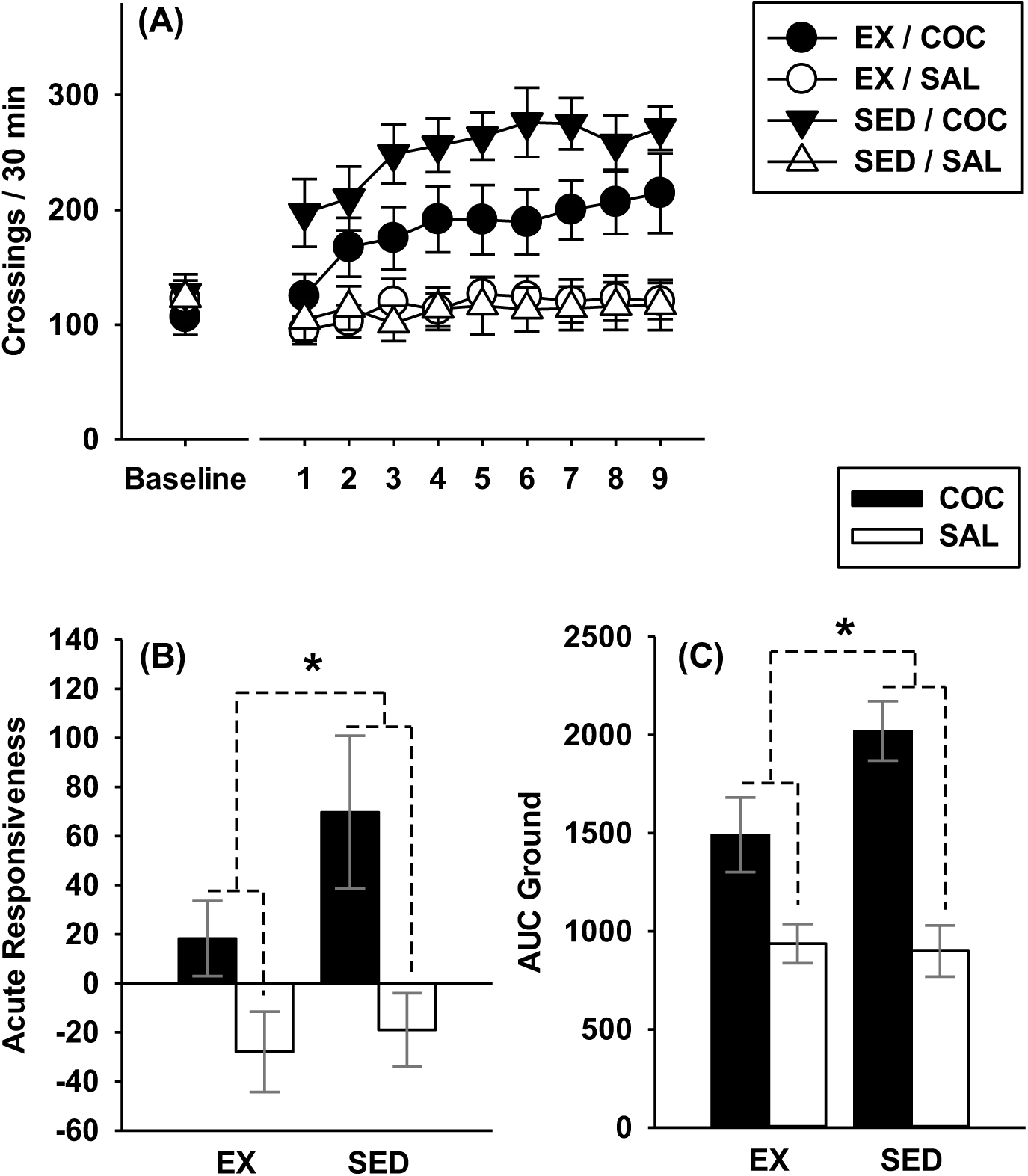
Acute responsiveness and initiation of sensitization. (A) Baseline locomotor activity (under saline) and initiation of locomotor sensitization over 9 once-daily sessions. (B) Acute responsiveness scored as the difference between values from the 1^st^ and baseline sessions. (C) Overall locomotor responsiveness over the initiation of sensitization scored as AUC ground. **\***significant interaction-related difference between the cocaine effect observed in exercised mice (EX/COC, n=24 vs EX/SAL, n=24) and that measured in sedentary mice (SED/COC, n=24 vs SED/SAL, n=24) taken at a threshold of 0.05. Bars represent 95% confidence intervals.

**Fig 4.**
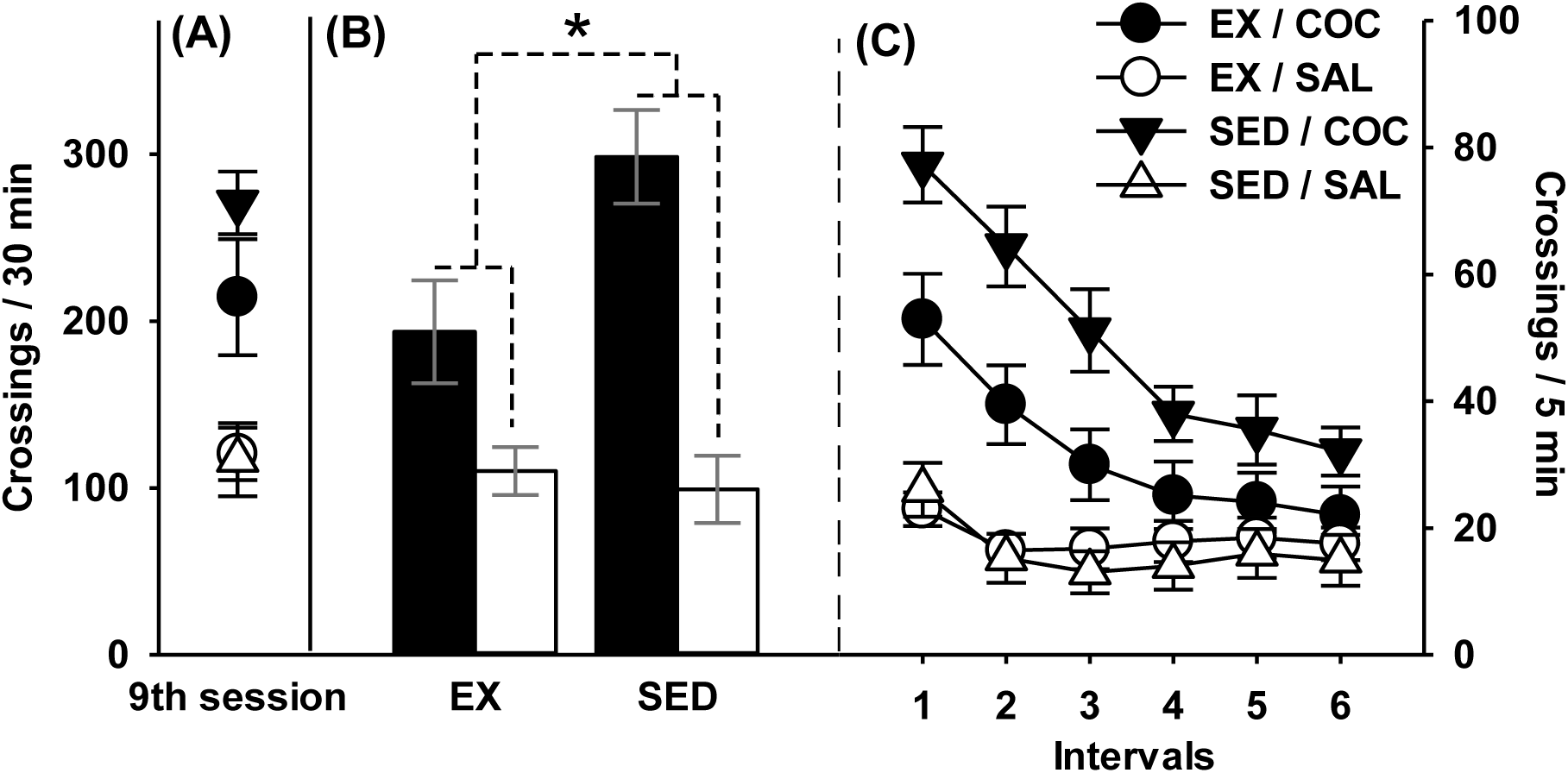
Long-term expression of sensitization. (A) Locomotor responsiveness on the last (9^th^) once-daily session (descriptive statistics). (B) Locomotor responsiveness on the test for expression of sensitization. *significant housing conditions × pharmacological treatment interaction: cocaine effect measured in sedentary mice (SED/COC, n=24 vs. SED/SAL, n=24) is greater than that observed in exercised mice (EX/COC, n=24 vs EX/SAL, n=24). (C) Time-course of locomotor responsiveness during the test for expression of sensitization (descriptive, no inferential statistics were conducted on these data). Bars represent 95% confidence intervals.

**Table 1.** Relationships between averaged pre-testing running distances and behavioral and neuro-imaging outcomes (Pearson correlation coefficient (*p*-values)).

|  | Acute Responsiveness | AUC ground | Expression of Sensitization | [<sup>18</sup>F]fallypride BP<sub>ND</sub> |
| --- | --- | --- | --- | --- |
| COC | -0.17 (0.43) | 0.43 (0.035) | 0.15 (0.50) | -0.24 (0.33)<sup>a</sup> |
| SAL | -0.11 (0.61) | -0.10 (0.66) | -0.03 (0.89) | -0.09 (0.75)<sup>b</sup> |

<sup>a</sup> n=19, <sup>b</sup> n=16, n=24 otherwise.

### [^18^F]fallypride microPET neuroimaging

Fig. 5 panel A displays representative [^18^F]fallypride BP_ND_ images of mice of the four groups. Panel B presents [^18^F]fallypride BP_ND_ measured in the striatum 24h after the expression of sensitization in exercised (EX/COC and EX/SAL) and sedentary (SED/COC and SED/SAL) mice. Due to technical problems (*i*.*e*. fails in proper intravenous radiotracer delivery at injection time), data from 23 mice (over 96) were not acquired or useable (EX/COC: n=5; EX/SAL: n=8; SED/COC: n=5; and SED/SAL: n=5). Panels C and D present the marginal means associated with main effects of housing conditions (EX: n=35; SED: n=38) and pharmacological treatment (COC: n=38; SAL: n=35) respectively. We found evidence for a moderate attenuating effect of aerobic exercise on [^18^F]fallypride BP_ND_ (*η*^*2*^*p* = 0.075, PS = 65, *t*_(50)_ = 2.01, *p* = 0.024). Additionally, cocaine-receiving mice exhibited higher [^18^F]fallypride BP_ND_ in striatum than their saline counterparts as supported by a large effect of pharmacological treatment (*η*^*2*^*p* = 0.170, PS = 74, *t*_(50)_ = 3.20, *p* = 0.001). However, crossed contrasts indicate that the interaction between housing conditions and the pharmacological treatment was clearly negligible (*η*^*2*^*p* < 0.005, *t*_(50)_ = 0.20, *p* = 0.42). Table 2 reports relationships between behavioral outcomes and [^18^F]fallypride BP_ND_. There was no evidence for association between behavioral outcomes and BP_ND_ values, with correlation coefficients *r*s varying from 0.005 to 0.37 (*p*-values ranging from 0.98 to 0.12).

**Table 2.**
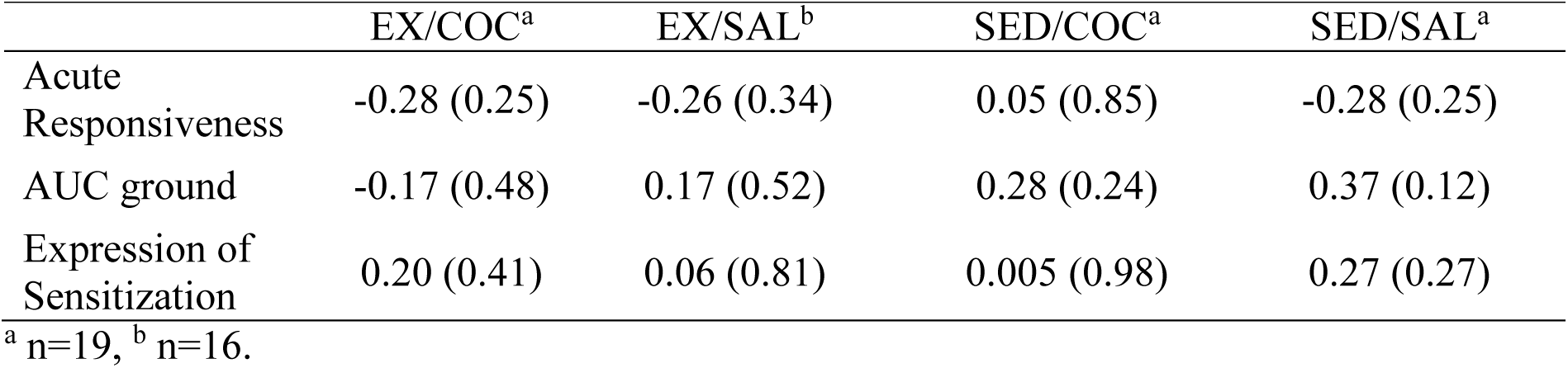
Relationships between behavioral outcomes and neuro-imaging outcomes expressed as [^18^F]fallypride BP_ND_ (Pearson correlation coefficients (p-values)).

|  | EX/COC<sup>a</sup> | EX/SAL<sup>b</sup> | SED/COC<sup>a</sup> | SED/SAL<sup>a</sup> |
| --- | --- | --- | --- | --- |
| Acute Responsiveness | -0.28 (0.25) | -0.26 (0.34) | 0.05 (0.85) | -0.28 (0.25) |
| AUC ground | -0.17 (0.48) | 0.17 (0.52) | 0.28 (0.24) | 0.37 (0.12) |
| Expression of Sensitization | 0.20 (0.41) | 0.06 (0.81) | 0.005 (0.98) | 0.27 (0.27) |

<sup>a</sup> n=19, <sup>b</sup> n=16.

**Fig 5.**
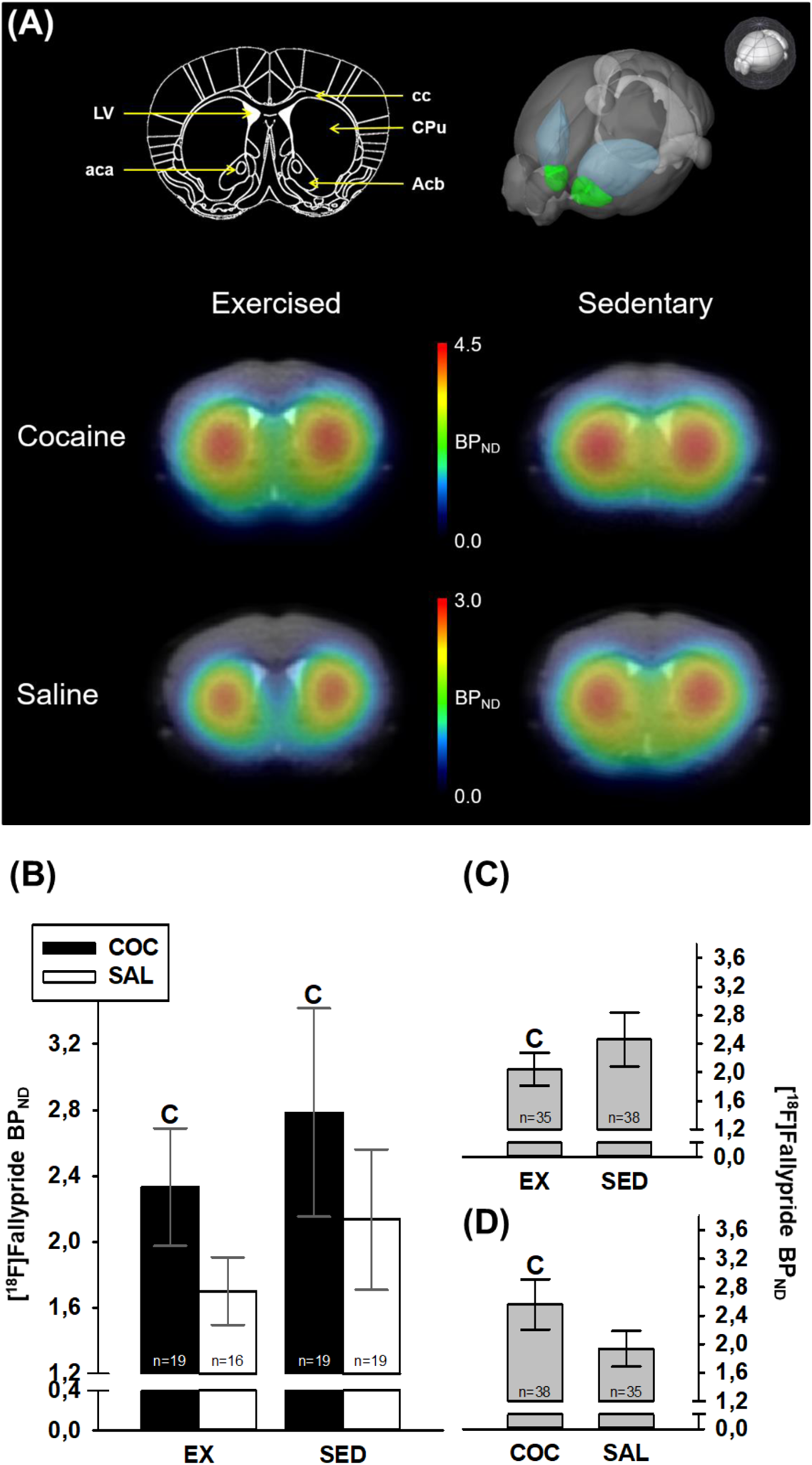
Neuroimaging outcomes. (A) Left upper panel: coronal slice image of the mouse brain at Bregma 0.8 mm, based on the mouse Atlas of Franklin and Paxinos^57^. (A) Right upper panel: 3D depiction of regions of interest showing the Caudate Putamen (in blue) and the Nucleus Accumbens (in green). (A) Bottom: representative [^18^F]fallypride BP_ND_ images of mice of the four groups, co-registered to their corresponding individual anatomical MRI. Note the difference in scales between the cocaine groups and the saline groups. (B) microPET-derived [^18^F]Fallypride BP_ND_ measured 24 h after expression of sensitization in exercised and sedentary mice. (C) Marginal means associated with the effect of housing conditions. (D) Marginal means associated with the effect of pharmacological treatment. **C**s indicate significant difference compared to the corresponding control group. Bars represent 95% confidence intervals.

## DISCUSSION

The main findings of the present study can be summarized as follows. (1) Previous results obtained in our laboratory were replicated by showing that free wheel running, a useful model for the study of voluntary aerobic exercise, induced preventive effects against acute and chronic locomotor responsiveness to a rewarding-like dose of cocaine (8 mg/kg) in female C57BL/6J mice. (2) The cocaine sensitized mice brain, at the time of long term expression, revealed a striatal increase in dopamine D2/3 receptors availability measured by [^18^F]fallypride microPET. (3) Voluntary wheel running was associated with attenuated dopamine D2/3 receptors availability measured by [^18^F]fallypride microPET.

Wheel-running, allowed throughout experimentation, mitigated acute locomotor responsiveness and both the initiation and the expression of locomotor sensitization to a representative (and hedonistic) dose of cocaine^46^. These results reproduce previous findings obtained in our laboratory^25,41^. This set of observations are consistent with those reported by Renteria Diaz and colleagues^24^, whose male Wistar rats continuously housed with a wheel for at least 5 weeks expressed little or no sensitized locomotor activity 15 days after 5 once-daily injections of 10 mg/kg cocaine. Smith and Witte^58^ also showed that continuous wheel-running exercise was effective at reducing the locomotor-activating effects of 3 and 10 mg/kg cocaine in Long-Evans females. Such attenuating effect was also observed in C57BL/6J mice housed in large home-cages comprising a running wheel as part of a composite housing environment made of inanimate objects and conspecifics^59,60^. More generally, our results add to extensive preclinical literature reporting preventive consequences of wheel-running exercise on behavioral markers of sensitivity to addictive properties of drugs of abuse^15^. Thus, the reliability of our model make it relevant for biological study of addiction, especially for the DA system which shows cross-species homologies between humans and well conserved neuroanatomy and circuit function^61^.

Physical exercise is known to act on DA system and to possess rewarding properties, like cocaine and other drugs of abuse^62,63^. For example, Greenwood and co-workers^63^ reported that Fischer344 rats exercising with a running wheel for 6 weeks preferred a compartment paired with the aftereffect of exercise (conditioning on alternate days) whereas no preference for the paired compartment appeared when the rats used the wheel for 2 weeks. They also showed that a 6-week continuous access to wheels resulted in increases in ΔFosB/FosB immunoreactivity in the nucleus accumbens (Acb), tyrosine hydroxylase (TH) mRNA levels in the ventral tegmental area (VTA) and delta opioid receptor mRNA levels in the Acb shell, whereas dopamine receptor D2 mRNA is reduced in the Acb core. It is thus tempting to ascribe the exercise-induced protective effects against the cocaine-induced locomotor sensitization (and its rewarding properties, since we used a rewarding dose of the drug) to its neuroplastic effects on dopaminergic neurotransmission.

To our knowledge, this is the first study reporting in vivo PET imaging of mouse striatum D2/3R under aerobic exercise in the context of cocaine sensitization. The first line of our imaging results suggests that the long term expression of cocaine sensitization is associated to an increase of D2/3R availability in the mouse striatum, as demonstrated by the large-sized main effect of cocaine on the [^18^F]fallypride BP_ND_. Those results are in line with studies suggesting that psychomotor sensitization to psychostimulant is linked to enhanced D2/3R availability^32,33,64^ which may explain the high locomotor response of psychostimulant sensitized mice to direct-acting D2 agonists^30^. Although previous reports, using either amphetamine or cocaine as well as nicotine, hypothesized that the behavioral sensitization could occur through an increase of the dopamine D2 high affinity state receptors availability (*i*.*e*. actively coupled to G-proteins)^38–40^, the [^18^F]fallypride radiotracer is not able to discriminate between the two affinity states of D2 receptors. Therefore our results seem not to rely on the proportion of high versus low affinity state but rather on the total amount of available receptors. Our work could be interpreted in light of the supposed enhanced activity of DA neurons in the VTA during cocaine sensitization^65^ where the increased D2/3R availability in the striatum may contribute to the enhanced locomotor response induced by repeated cocaine injections.

The most innovative outcome of our study is the decreased D2/3R availability in the striatum of exercised mice, compare to their sedentary counterparts as highlighted by the medium-sized main effect of the housing condition on the [^18^F]fallypride BP_ND_. This is in line with previously reported results were wheel-running has been shown to provoke a reduction of D2R mRNA in Fisher 344 rats Acb core^63^. The authors also reported an increased TH mRNA level in the VTA leading them to propose that voluntary exercise can increase the synthetic capacity of DA in the striatum^63,66^. It seems that these modifications could be related to neuroplastic events induced by an enhanced expression of ΔFosB transcription factor in the Acb nucleus^63^. Interestingly, a sustained accumulation of ΔFosB in the Acb nucleus was also described following chronic cocaine exposure, as well as for others drugs of abuse^67–69^. This common feature led Greenwood and coll. to the assumption that neuroplasticity induced by voluntary exercise could alter DA neurotransmission in the mesolimbic reward pathway which may contribute then to the beneficial effects of exercise on cocaine sensitization^63^. This is supported by data reporting that chronically exercised Sprague Dawley rats displayed a lower DA release, and a hampered DA reuptake, under amphetamine challenge^70^. Although, our work revealed the absence of any significant interaction between the housing conditions and the pharmacological treatment, meaning that there was no specific biological action of free wheel running on cocaine-sensitized mice (cf. previous paragraph), our data add evidences to the previously mention hypothesis. Moreover, physical exercise has been shown to reduce the basal level of DA in the rat striatum, which accord to the hypothesis of a reduced DA tone resulting from chronic physical exercise despite an enhanced synthetic capacity^71^. We suggest that voluntary physical exercise acts on DA mesolimbic reward pathway partly through a reduced D2/3R availability in the mouse striatum, which trait makes exercised mice more resilient to psychomotor effects of cocaine.

Some limitations of this study warrant mention, in particular those related to intrinsic limitations of in vivo microPET imaging in mice. First of all, the [^18^]Fallypride radiotracer displays a nearly equal affinity for D2 and D3 dopamine receptors in vivo^72^ and cannot therefore distinguish between them. Thus, [^18^]Fallypride BP_ND_ parameter primarily reflect a combination of signals from D2 and D3 receptors. It has been suggested that up to 20% of the [^18^]Fallypride binding may be due to D3R in vivo^73^. This is of relevance considering that D3R are highly expressed in the mesolimbic DA system and involved in the pathophysiology of addiction^8^. Strikingly, their expression seemed to be increased in nicotine behavioral sensitization^74^. Determining the respective proportion of D2R and D3R in our present results would definitely need further investigations. The second technical limitation in in vivo microPET investigation is related to the spatial resolution of the microPET scanner, which is about 1.5 mm^2^ in our setup (*i*.*e*. Siemens Focus 120)^49^. This has of course to be taken into account when attempting to achieve in vivo molecular imaging of mouse brain. The mouse brain atlas implemented in Pmod 3.7, derived from the work of Mirrione and coll., didn’t allow to analyze separately the ventral and the dorsal parts of the striatum, the Accumbens nucleus (Acb) and the Caute-Putamen (CPu) respectively^51^. As a consequence the ROI analyzed to compute the [^18^F]fallypride BP_ND_ is constituted by the entire striatum. This is of interest as the DA inputs within the Acb nucleus hail from the Ventral Tegmental Area (VTA) whereas the DA afferences throughout the CPu originate from the Substancia Nigra (SN). On one hand, DA neurons of the VTA are known to play a major role into the reward and motivation processes^75^, and to their pathological counterpart that is addiction. On the other hand, DA neurons lying in the SN are controlling motor functions. Intuitively, one would think that the [^18^F]fallypride BP_ND_ modifications we are describing here are primarily located in the Acb nucleus. However, D2R are involved in locomotor activity and may be implicated in [^18^F]fallypride BP_ND_ variations induced by physical exercise^76^. Our results may be conflicting with those of Vuckovic and coll., reporting an increased [^18^F]fallypride BP_ND_ in a mouse model relevant for the study of Parkinson’s disease under exercise condition. However, the pathophysiological paradigm shift prevents any direct comparison of the data, as the increase in the [^18^F]fallypride BP_ND_ there were reporting was seen in the DA terminals depleted group, whereas they report no differences between saline and saline plus exercise mice^76^.

Clinical assessment of D2/3R in methamphetamine users under behavioral intervention with exercise training has been achieved using [^18^F]fallypride PET imaging^77^. Interestingly, they reported an increased [^18^F]fallypride BP_ND_ in the whole striatum of patients include into the exercise group in comparison to those of the control group (*i*.*e*. methamphetamine users under behavioral intervention with educational training). Thus, the author suggest that under depleted striatal D2/3R availability, as shown by the methamphetamine users, physical exercise may increase the availability of these receptors. However, they tempered themselves their results given the relative small size of the samples. Besides, the absence of healthy control in the study design precludes conclusions drawing on the effect of physical exercise on striatal D2/3R availability in physiological conditions. Moreover, we stress here that our results focused on the sensitization process (and the associated long term expression) which is thought to be a useful model to investigate early events in the natural history of addiction pathophysiology (*e*.*g*. recreational use). On the contrary, the work of Robertson and coll. included patients under DSM-IV-TR criteria for methamphetamine dependence, meaning that they already have a long history of methamphetamine abuse. Furthermore, the included patients are under complete withdrawal which could impact by itself DA neurochemical processes and makes a clear difference with our study design.

In conclusion, we report a replication study of a protective effect of wheel-running exercise on both induction and expression of cocaine locomotor sensitization in female C57/BL6J mice. Our findings show that exercise-induced neuroplasticity within mesolimbic DA pathway include a reduced D2/3R availability in the striatum, while cocaine locomotor sensitization is associated with an increased D2/3R availability in this brain area. While, further investigations are required to unravel molecular mechanisms by which exercise affects dopaminergic signaling, our work contributes to the neurobiological understanding of physical exercise and its positive impact on addictive behaviors.

## ACKNOWLEDGEMENTS

The present work was supported by grants from the “Fonds National de la Recherche Scientifique” (FNRS) and the University of Liège “Fonds Spéciaux pour la Recherche” obtained by Ezio TIRELLI and Alain PLENEVAUX.

## AUTHOR CONTRIBUTIONS

G.B. contributed to the conception of the study, acquired and analyzed the imaging data, interpreted the data and wrote the paper. L-F.L contributed to the design of the study, acquired and analyzed the behavioral data, interpreted the data and wrote the paper. M.A.B. contributed to the acquisition and analysis of the imaging data and wrote the paper. M.E.S. contributed to the acquisition and analysis of the imaging data. C.L. performed all the radiosyntheses and have drafted the paper. A.L. contributed to the conception of the study and substantially revised the work. A.P. and E.T. have funded this work, contributed to the conception of the study, interpreted the data and substantially revised the work.

